# Science abhors a surveillance vacuum: detection of ticks and tick-borne bacteria in southern New Mexico through passive surveillance

**DOI:** 10.1101/2023.09.25.559416

**Authors:** Paige R. Harman, Nicole L. Mendell, Maysee M. Harman, Puck A. Draney, Anna T. Boyle, Matthew E. Gompper, Teri J. Orr, Donald H. Bouyer, Pete D. Teel, Kathryn A. Hanley

**Author notes:** Department of Biology, Wesleyan University, Middletown, Connecticut, United States of America (Maysee M. Harman).

## Abstract

Robust tick surveillance enhances diagnosis and prevention of tick-borne pathogens, yet surveillance efforts in the U.S. are highly uneven, resulting in large surveillance vacuums, one of which spans the state of New Mexico. As part of a larger effort to fill this vacuum, we conducted both active and passive tick sampling in New Mexico, focusing on the southern portion of the state. We conducted active tick sampling using dragging and CO_2_ trapping at 45 sites across Hidalgo, Doña Ana, Otero, and Eddy counties periodically between June 2021 and August 2022. We also conducted opportunistic, passive tick sampling in 2021 and 2022 from animals harvested by hunters or captured or collected by researchers and animals housed in animal hospitals, shelters, and farms. All pools of ticks were screened for *Rickettsia rickettsii, R. parkeri, R. amblyommatis, Ehrlichia ewingii*, and *E. chaffeensis*. Active sampling yielded no ticks. Passive sampling yielded 497 ticks comprising *Carios kelleyi* from pallid bats, *Rhipicephalus sanguineus* from dogs, mule deer, and Rocky Mountain elk, *Otobius megnini* from dogs, cats, horses, and Coues deer, *Dermacentor parumapertus* from dogs and black-tailed jackrabbits, *D. albipictus* from domesticated cats, mule deer and Rocky Mountain elk, and *Dermacentor spp*. from American black bear, Rocky Mountain elk, and mule deer. One pool of *D. parumapterus* from a black-tailed jackrabbit in Luna County tested positive for *R. parkeri*, an agent of spotted fever rickettsiosis. Additionally, a spotted fever group *Rickettsia* was detected in 6 of 7 *Carios kelleyi* pools. Two ticks showed morphological abnormalities; however, these samples did not test positive for any of the target pathogens, and the cause of the abnormalities is unknown. Passive surveillance yielded five identified species of ticks from three domestic and six wild mammal species. One tick pool from a black-tailed jackrabbit was found to harbor *Rickettsia parkeri*, and six pools of *Carios kelleyi* ticks, argasid ticks that have been reported to bite humans, were found to harbor a spotted fever group *Rickettsia*. Our findings update tick distributions and inform the public, medical, and veterinary communities of the potential tick-borne pathogens present in southern New Mexico.

## Introduction

Ticks are vectors for a wide range of pathogens, including bacteria, parasites, and viruses [1]. Lyme disease (*Borrelia burgdorferi*) is the most common vector-borne disease in humans in the U.S., and a variety of other tick-borne bacteria, parasites and viruses pose a serious threat within the country, including *Anaplasma phagocytophilum* (anaplasmosis), *Rickettsia rickettsii* (Rocky Mountain spotted fever), *Babesia microti* (babesiosis), and Powassan virus [2, 3, 4]. Timely diagnosis of tick-borne diseases by health care providers, use of appropriate personal protection by the public, and recognition of increased incidence or expansion of ticks and tick-borne pathogens by the scientific and public health communities depend upon comprehensive and sustained tick surveillance [5, 6]. Unfortunately, tick surveillance has been extremely uneven across the US, leaving dangerous surveillance vacuums [7, 8].

One of the most extensive of these surveillance vacuums occurs in the state of New Mexico. *Ornithodoros turicata*, the vector of relapsing fever, was first reported in New Mexico in 1908 [9]. The first records of tick-borne relapsing fever were reported in Bear Creek Canyon, Jefferson County, Colorado in 1915 and 1918 [9]. After this, relapsing fever expanded rapidly throughout the U.S., and was first detected in New Mexico in 1934 [9]. In the century since this discovery, there has been little surveillance of ticks in New Mexico, and most studies have focused on a limited number of hosts or a limited geographic area. In 1952, Allen and Kennedy first reported *Otobius megnini* and *Dermacentor albipictus* from a mountain sheep in south-central New Mexico [10], and the former species has been detected in ungulates in the state in several studies since [11]. In 1984 Eads and Campos provided a brief synopsis of human parasitism by *O. megnini* in the state, indicating that parasitism by these ticks occurs on humans, albeit infrequently, thereby exposing them as a larger public health concern than previously understood [12]. In 1975, Meleney discovered *O. megnini* and *Dermacentor albipictus* on collared peccaries in southwestern New Mexico [13]. Between 1984 to 1988, working in eastern New Mexico, Pfaffenberger and colleagues detected one *O. turicata* on a prairie dog [14], *Dermacentor parumapertus* on rodents and rabbits [15], and *Ornithodoros parkeri* and *Haemaphysalis leporispalustris* on rabbits [16]. In 1996, Cokendolpher and Polyak discovered *Ixodes conepati* in a cave in eastern New Mexico [17]. In 2001, Steinlein et al. detected *Carios* (*Ornithodoros) kelleyi* and *O. rossi* on a variety of bat species, constituting the first report of these species in New Mexico [18]. In 2006, James et al. summarized distribution of *Dermacentor andersoni* across the U.S.; this species was found attached to a wide array of both wild and domestic mammals as well as humans in northern New Mexico [19]. In 2020, Hecht et al. investigated the occurrence of *Amblyomma maculatum* sensu lato ticks and associated *Rickettsia parkeri* bacteria in far southwestern New Mexico and reported the first instance of *Amblyomma maculatum* Koch sensu lato (s.l.) infected with *R. parkeri* [20]. Most recently, in 2023, Grant et al. assessed the distributions and lineages of temperate and tropical *Rhipicephalus sanguineus* sensu lato within the U.S. and found that the *R. sanguineus* sensu lato in New Mexico belong to the temperate lineage [21]. Even accounting for the fact that our search terms (New Mexico combined with tick or ectoparasite) in Pubmed and Google Scholar likely failed to return some relevant studies, nonetheless these publications represent a relatively small surveillance effort in the fifth largest state, in land area, in the U.S.

In the current study, we conducted a broad effort in active and passive surveillance of ticks in New Mexico (Figure 1), focusing predominantly on the south-central and southwestern region of the state, close to the US-Mexico border. South-central and southwestern New Mexico span a large, arid region encompassing the Chihuahuan desert, Montane, High Plains and Madrean Archipelago ecoregions [22]. Due to this landscape diversity, this region represents a hotspot of vertebrate diversity [23], providing a plethora of host species for tick attachment. Additionally, the state is home to an established ranching and farming enterprise providing abundant horses, cattle, sheep and goats as tick hosts. Both wild and domestic animals may move over substantial distances in search of food and water, offering great potential for tick dispersal. However, the extreme daily fluctuations in temperature and general aridity of the region may impede direct tick movement and decrease survival of eggs and free-living ticks [24-30]. Conversely, some species like *R. sanguineus* are xerophilic, some nidicolous species may find refuge from climatic extremes in animal burrows, and some bacterial pathogens can enhance tick host desiccation resistance and cold tolerance [24-30].

**Fig 1.**
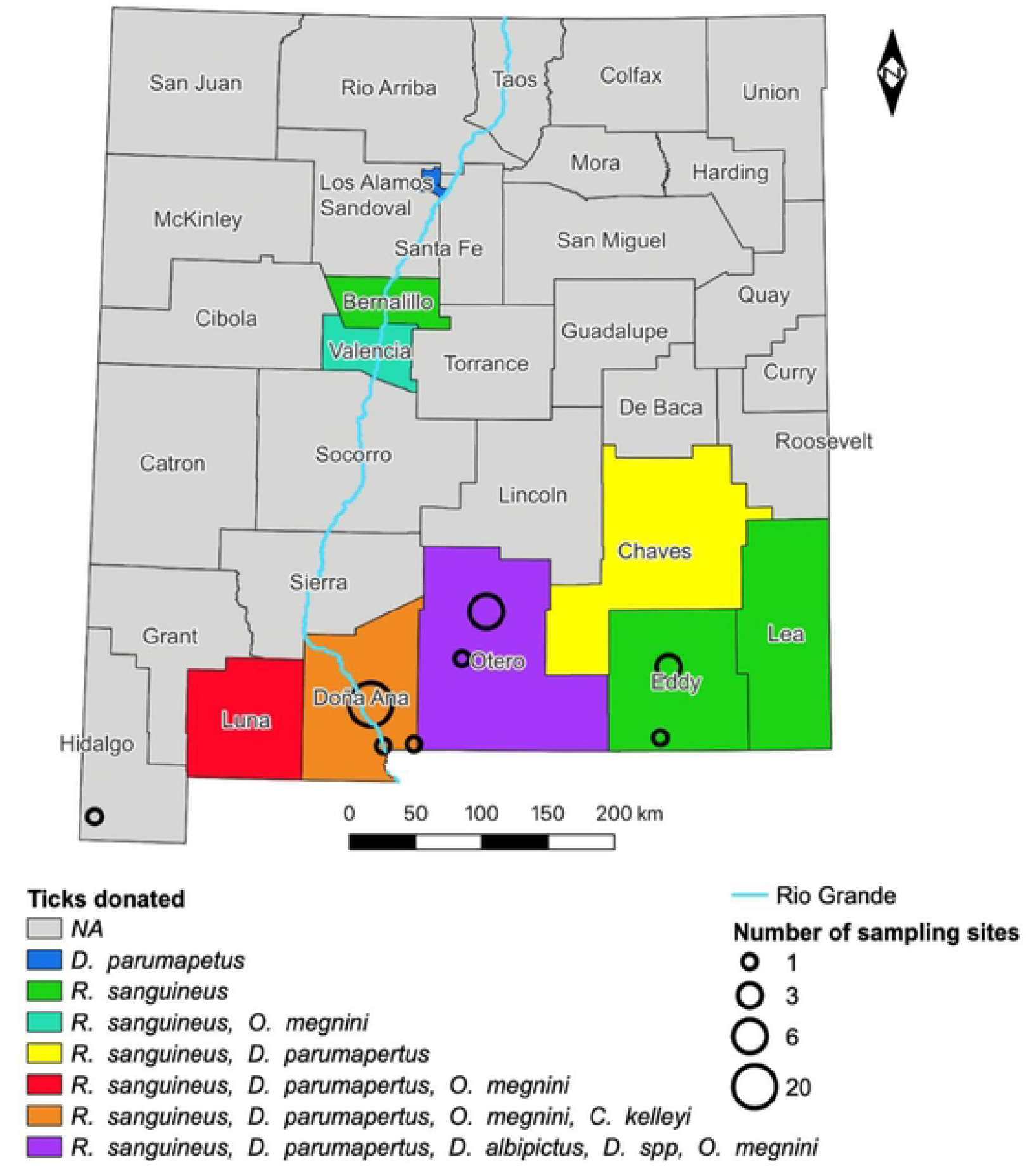
Map representing locations of both active (open circles) and passive (shaded counties) tick sampling. Circle diameter indicates number of sampling sites, colors indicate tick species collected from each county as listed above.

While active surveillance across multiple New Mexico counties yielded no ticks, passive surveillance of a variety of wildlife and domestic mammals yielded almost 500 ticks, comprising five identified species. We found molecular evidence from one tick pool for the presence of the pathogenic bacteria, *R. parkeri*, which causes human rickettsioses, as well as evidence for a minimally characterized spotted fever group *Rickettsia*. To our knowledge, our findings expand the documented range of the ticks *D. albipictus, O. megnini*, and *C. kelleyi* and adds to the documentation of rickettsial species in argasid tick.

## Materials and Methods

### Active Sampling

From June 2021 to August 2022, active tick surveillance was conducted at sampling sites across southern New Mexico at sites in Hidalgo, Doña Ana, Otero, and Eddy counties (**Fig 1** and **Table S2**). Due to extreme fluctuations in temperature and rainfall across the state, climate averages have little meaning, but generally the southern deserts receive less than 25 cm of rain annually, most of which falls during the summer monsoons [31]. June and July are the hottest months, with temperatures regularly exceeding 37°C. Collection sites (**Fig 1**) were chosen based on accessibility and encompassed a range of elevation and habitat. We included one location in southwestern NM where *A. maculatum* had previously been collected [20]. Study areas ranged in elevation from 1,000 m to 2,810 m.

Active sampling included flannel cloth drags and resting CO_2_ traps [32]. At each sampling site, one tick trap was placed and two drags were performed. Tick drags were conducted in the morning or in the evening between 06:30 and 12:00 or 14:30 and 20:30. The drags consist of a white flannel sheet (1.2 m × 1.2 m), a wooden dowel, and rope (2 m). Drags were pulled slowly along the ground in a straight line either East to West or North to South for a distance of 100 m, checking for ticks on both sides of the cloth every 10 m. CO_2_ traps consisted of a white flannel sheet (1.2 m × 1.2 m) on which was placed a ½ gallon igloo drinking cooler with eight 1 mm holes containing 1 kg of dry ice. CO_2_ traps were placed in the same time windows that dragging was conducted and left out for 3 hours or overnight before collection, as specified in **Table S2**.

### Passive Sampling

A network of local institutions and collaborators who were removing ticks from live animals in the course of normal care (veterinarians, pet daycares, animal shelters and farmers/ranchers) contributed ticks. New Mexico State University researchers who were capturing or collecting animals also contributed ticks; these included all samples from jackrabbits (NMSU IACUC 2020-019), bats (NMSU IACUC 2020-005) and bears (NMSU IACUC 2021-013). Additionally, local hunters were asked to allow NMSU personnel to screen harvested animals for ticks, which sometimes consisted of whole carcasses and sometimes only partial carcasses, at NM Department of Game and Fish (NMDF) chronic wasting disease check stations. Tick contributors were provided with a tick collection kit consisting of a consent and instruction form, datasheet for animal information (host type, county, date, host habitat, host age, number of ticks, location on body tick was collected), and 1.5 ml snap cap microcentrifuge tubes containing 1 ml 70% ethanol. Ticks collected from these sampling methods were either stored in 70% ethanol at a maximum of 10 ticks per pool or moved immediately to a -80°C freezer or maintained as live specimens in plastic bags with damp paper towels. All ticks were initially identified using morphological characteristics following Lado et al. [33] in the Hanley lab at New Mexico State University, and then shipped to the Bouyer lab at University of Texas Medical Branch for confirmation of identification and screening for bacterial pathogens.

### DNA Extraction

Live specimens were surface decontaminated with a 3% bleach solution, followed by a 70% ethanol solution and then rinsed with sterile water. Ticks were pooled based on species, host, sex, and stage with a maximum of 10 adult ticks or 25 nymphs per pool. Ticks received alive or with uncompromised cold chain were sagittally bisected, with half used for screening and the second half stored at -80°C for future analysis. Pools were homogenized with a TissueLyser II (Qiagen, Germantown, MD) for 3 min at 28 Hz in 2 mL microcentrifuge tubes in 1X sterile PBS or four volumes of DNA/RNA Shield (Zymo Research, Irvine, CA) and two 5 mm stainless steel grinding balls. Following homogenization DNA was extracted from the pools using either a Qiagen DNeasy blood and tissue kit (Germantown, Maryland) using the manufacturer’s instructions or co-extraction of DNA and RNA was accomplished with a Quick DNA/RNA MiniPrep Plus Kit (Zymo Research, Irvine, CA). The quality and quantity of DNA and RNA were analyzed by either NanoDrop 1000 or NanoDrop One (Thermo Fisher Scientific, Wilmington, DE) prior to pathogen detection. Morphological identification of *Carios kelleyi* and nymphal ticks were confirmed through analysis of the mitochondrial 16s rDNA sequence by PCR amplification using primers 16S+1 and 16S-1 (**Table 1**) [34-37].

**Table 1.**
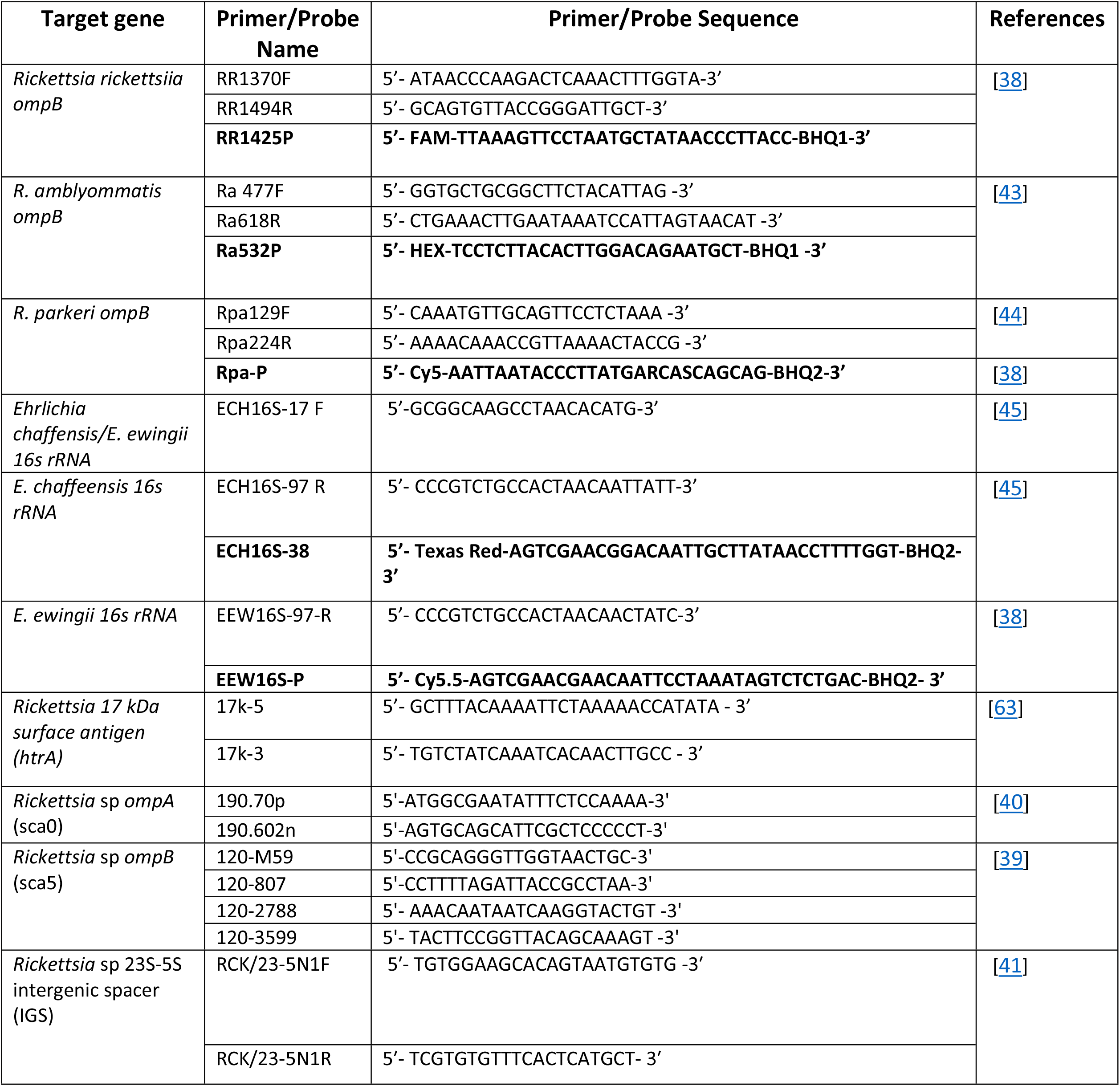
List of primers and probes (bold font) used for pathogen detection in tick samples.

### Pathogen Screening

Tick pool DNA was screened by multiplex for bacterial presence using a multiplex for *Rickettsia rickettsii, R. parkeri, R. amblyommatis, Ehrlichia ewingii*, and *E. chaffeensis* using iQ Multiplex Powermix (Bio-rad, Hercules, CA) in 20 μL reactions with 2 μL of DNA extract [38] (**Table 1**). Plasmids containing a single copy of each targeted gene were employed as positive controls.

Positives identified by the multiplex screening assay were confirmed by additional sequencing of rickettsial htrA (17 kDa) 17k3 and 17 k5, Sca0 (ompA); 190.70p and 190.602n, Sca5 (ompB);120-M59 and 120-807 and/or 120-2788 and 120-3599, or 23S-5S IGS; RCK/23-5N1F and RCK/23-5N1R (**Table 1**) [39-41]. DNA extracted from *R. sibirica* culture was utilized as a positive control. Reaction mixes for PCR were prepared in 25 μL reactions using 5PRIME HotMasterMix (Quantabio, Beverly, MA) and 800 μg/mL bovine serum albumin (BSA) water [42]. Previously published PCR thermal cycling conditions were used (**Table 1**) and resultant products were separated and visualized on an 1.5% agarose gel stained with ethidium bromide.

### Nucleotide Sequencing and Phylogenetic Analysis

Amplicons were purified with ExoSAP-IT (Applied Biosystems, Waltham, MA) and sequenced using a 3130/3130xl Genetic Analyzer (Applied Biosystems, Waltham, MA) according to manufacturer’s protocol. Resulting sequences were aligned using Lasergene 17 Seqman Pro software (DNASTAR, Madison, WI) and compared with published gene sequences available in GenBank using the Nucleotide Basic Local Alignment Search Tool-BLASTn (https://blast.ncbi.nlm.nih.gov/Blast.cgi). Representative sequences for htrA, ompA, and ompB from 16 rickettsial strains with complete genomes were downloaded from GenBank. Sequences were concatenated and aligned in MEGA 11 software with Clustal W and the final alignment presented 1887 nucleotides.

To investigate *Rickettsia* sequences from *C. kelleyi* ticks, phylogenetic analysis was performed using the Maximum Composite Likelihood approach with the General Time Reversible substitution model and gamma distribution. Branch supports were generated by bootstrap (1000 replicates). The tree was rooted with *R. akari* species from the transitional group.

## Results

### Active Sampling

Active sampling at 45 sampling sites (**Supporting Information: Table S2**) distributed across southern New Mexico did not yield any tick specimens.

### Passive Sampling

From June 2021 to August 2022, 497 ticks comprising the species *Carios kelleyi, Rhipicephalus sanguineus, Dermacentor albipictus, Dermacentor parumapterus* and *Otobius megnini* were collected by tick donators from 3 different species of domestic mammals and 6 different species of wild mammals (**Fig 2** and **Table S1**). Twenty-five of the tick pools collected were acquired from unknown hosts as a result of incompletely filled datasheets from tick donators. The location of one adult tick was listed as Gaines County, Texas; presumably this individual was visiting New Mexico.

**Fig 2.**
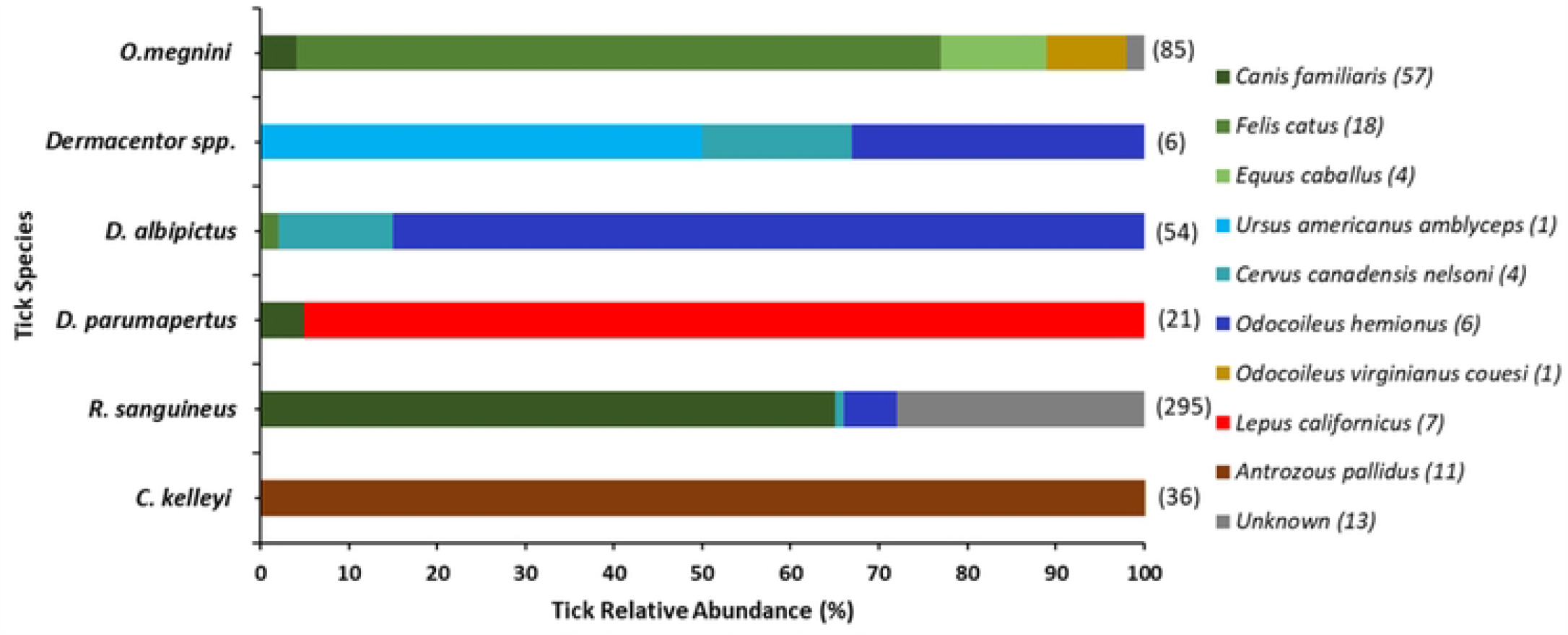
Relative abundance of tick species collected per host during passive sampling from June 2021 to October 2022 in New Mexico. Colors indicate host species from which ticks were taken, with the number of individual hosts in parentheses to the right of host name. Common names for each species are dog (*Canis familiaris*), cat (*Felis catus*), horse (*Equus caballus*), American black bear (*Ursus americanus amblyceps*), elk (*Cervus canadensis nelson*), mule deer (*Odocoileus hemionus*), Coues deer (*Odocoileus virginianus couesi*), black-tailed jack rabbit (*Lepus californicus*) and pallid bat (*Antrozous pallidus*).

Of the 180 tick pools screened, one pool of *D. parumapertus* from a black-tailed jackrabbit (*Lepus californicus*) in Luna County was positive for *R. parkeri* (**Table 2**), and obtained sequences were deposited in GenBank with accession numbers OR168635, OR168636, and OR168637. Sequence analysis in BLASTn showed 99.43% identity for *ompA* (493bp) and 99.74% (784bp) and 100% (712bp) for portions of the *ompB* to *R. parkeri* strain Black Gap.

**Table 2.** Screening results for one pool of *D. parumapertus* ticks from a black-tailed jackrabbit (*Lepus californicus*).

| Tick Species | No. ticks in Pool | Tick Stage & Sex | NM County | *Rickettsia* |  |  | *Ehrlichia* |  |
| --- | --- | --- | --- | --- | --- | --- | --- | --- |
|  |  |  |  | *parkeri* | *amblyommatis* | *rickettsia* | *chaffeensis* | *ewingii* |
| *Dermacentor parumapertus* | 3 | Adult F | Luna | + | - | - | - | - |

A rickettsial species was detected in six out of seven pools of *Carios kelleyi* ticks collected from pallid bats (*Antrozous pallidus*). Sequence analysis in BLASTn showed 99.43% identity for *htrA* (530bp) to *Rickettsia rickettsii*, 100% for *ompA* (494bp) to *Rickettsia* endosymbiont of *Carios kelleyi*, and 98.99% for *ompB* (760bp) to *R. peacockii*. Representative sequences were deposited in GenBank with accession numbers OR168632 OR168633, and OR168634 (**Fig 3**). Due to the conservation of the *ompB* gene sequence between *R. rickettsii* and the Rickettsial species identified in this study and unavailability of this minimally characterized bacterial sequence in Genbank, cross-reaction was observed in the species specific rtPCR assay.

**Fig 3.**
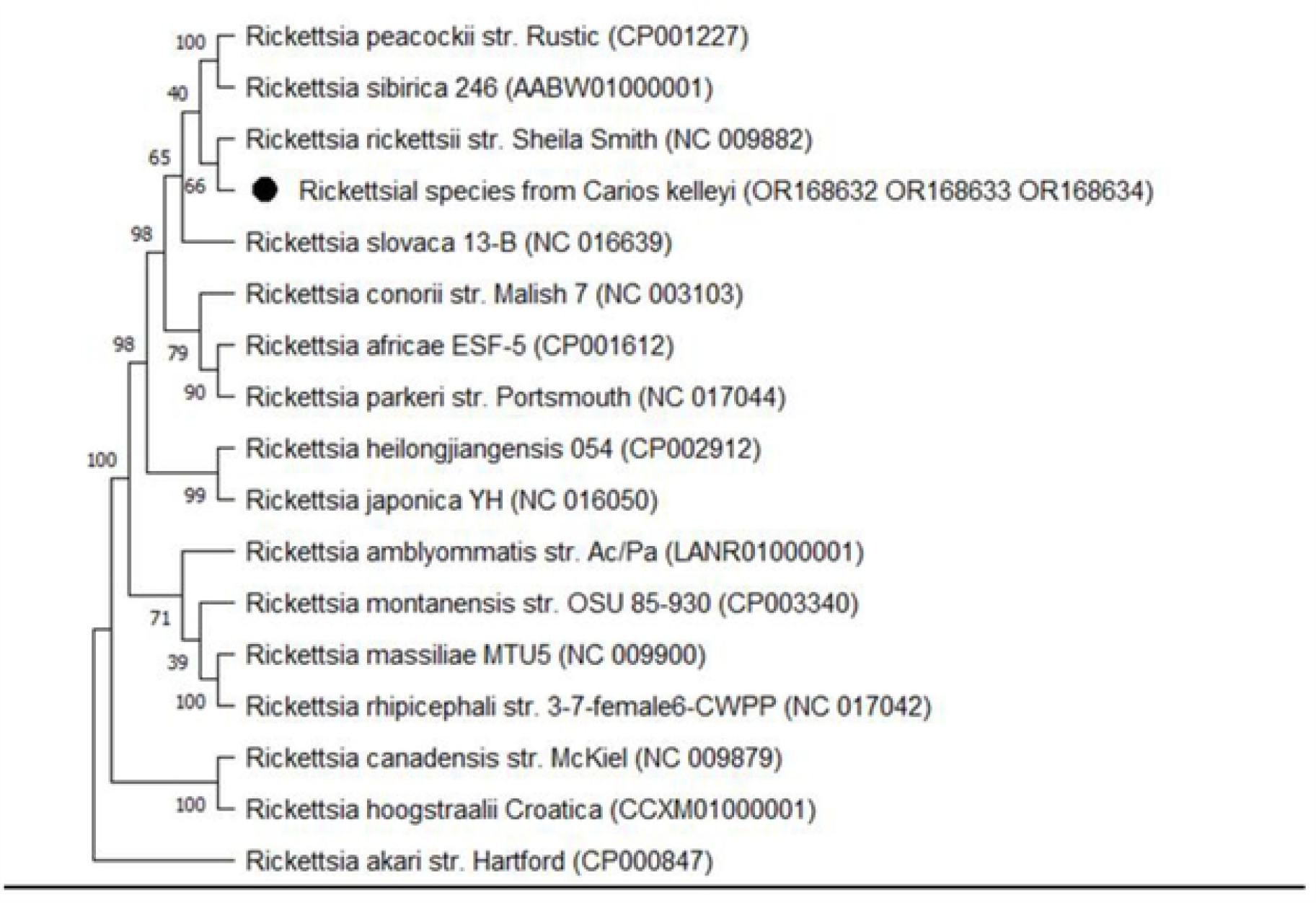
Consensus phylogenetic tree of the concatenated *htrA, ompA*, and *ompB* genes for the sequences obtained from the *Rickettsial* species detected in *Carios kelleyi* ticks indicated by the black dot.

### Morphological Abnormalities

Two *Rhipicephalus sanguineus* ticks exhibited morphological anomalies. Both samples were engorged adult females collected from a single dog. Both specimens showed abnormalities in the cuticle and throughout the alloscutum (**Fig 4**). The vesicular structures are observed throughout the tick bodies, where the coxa connects to the idiosoma, along where the scutum ends and the alloscutum starts, and around the spiracular plate. These abnormalities were not detected on any other tick samples.

**Fig 4.**
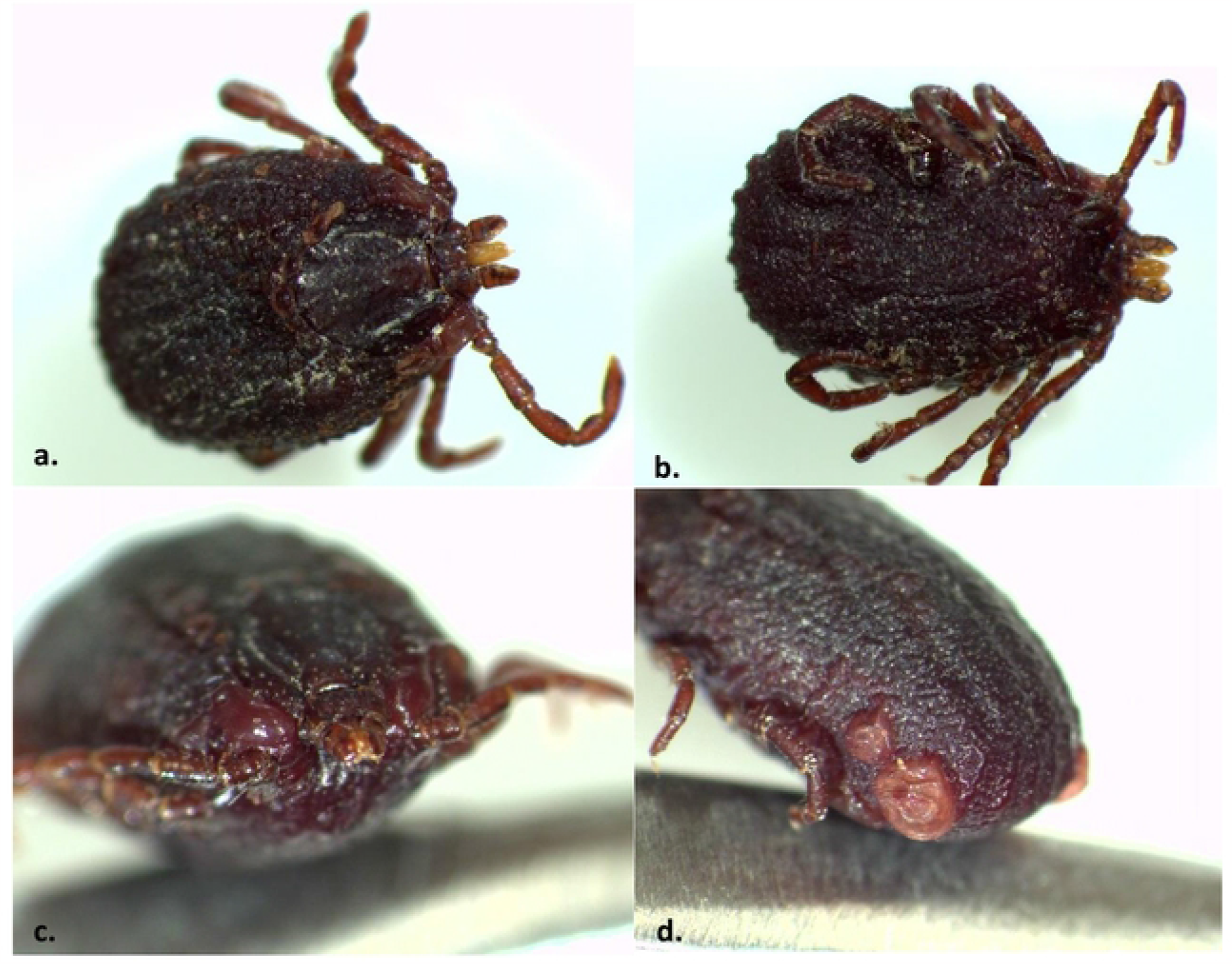
Two *R. sanguineus* engorged females (a, b) with anomalies around the coxa, throughout the alloscutum and along the spiracular plate (c,d).

## Discussion

Our passive surveillance efforts update and extend the known range of one tick species, *C. kelleyi*, within New Mexico. Given that ticks were obtained directly from hosts, we did not attempt to narrow the site of collection beyond county. No new tick-host associations were discovered. However, the attachment of *D. albipictus*, also known as the “winter tick”, to cats (*Felis catus*) is notable as this tick species is commonly associated with larger hosts [46]. Another association of interest is the discovery of *O. megnini* on cats, as these ticks are a one host ectoparasite which are usually found on large ungulates and have been documented to bite humans [47]. Our findings indicate cats offer a potential route for transport of these ticks into homes.

Additionally, we have demonstrated a broader vector range of tick-borne pathogens in New Mexico than formerly known. *Rickettsia parkeri*, and another *Rickettsia* agent were detected within our tick samples. While *R. parkeri* has been reported in New Mexico from *A. maculatum* sensu lato, this is the first report of *R. parkeri* from *Dermacentor spp*. in New Mexico [20]. *Rickettsia parkeri* Black Gap was characterized from a *D. parumapertus* tick isolate collected in Brewster County, Texas in 2015 and closely related bacteria have been identified in the same tick species from Utah and Arizona in the Western United States and the states of Sonora and Chihuahua in Northern Mexico [48, 49]. Moreover, *R. rickettsii, Ehrlichia canis, Anaplasma phagocytophilum* and *A. platys* have previously been identified in ticks (*R. sanguineus*) and dogs in Ciudad Juarez, Mexico, a city 40 km from the southern border of New Mexico [50], suggesting that further surveillance may also have yielded these pathogens as well. Molecular evidence of rickettsial species from *Carios kelleyi* ticks has been reported in Iowa and Kansas [51]. Our detection of a spotted fever group *Rickettsia* in *C. kelleyi* ticks from pallid bats, which often roost in or near human-made structures, merits further investigation.

Despite extensive efforts, active sampling in this study yielded no ticks, even in counties where passive sampling was successful. This scarcity contrasts sharply with active sampling in more mesic regions, which often yields, on average, one to ten ticks per 100 m drag [52-54]. Clearly ticks are surviving in the high desert and mountain habitats of our region, but where they occur when they are not attached to hosts remains unknown. It is possible that ticks occur at the surface at such low densities that our active sample efforts were insufficient to detect them. Seasonal variation may also play a role in sampling success: at the same sites in southwest New Mexico, we collected no ticks in April, during the dry season, while Hecht et al. [20] collected *A. maculatum* in August, during the monsoon season. Alternatively, ticks may be concentrated in refuges such as animal burrows, nests, caves, and other pits [9] that were outside the range of our drags and traps. More study and alternative sampling methods are needed to delineate the climatic ranges of ticks in the desert southwest and the strategies that they use to survive in a region that appears highly inhospitable for this taxon.

A handful of studies have reported results of surveillance for tick-borne pathogens in humans or wildlife in or near southern New Mexico. In 2002, eastern Arizona experienced an outbreak of Rocky Mountain spotted fever of epidemic proportions, intensifying public awareness and concern of the risk of tick-borne disease in the region [55]. Other studies within the desert southwest investigating the desert cottontail (*Sylvilagus audubonii*), American black bear (*Ursus americanus*), mule deer (*Odocoileus hemionus*) and coyote (*Canis latrans)* have detected the tick-borne pathogens *F. tularensis*, a novel *Borrelia spp*., *R. parkeri* and *Rickettsia spp*., underscoring the need for continued investigation [55-58]. Mule deer and American black bears were also included in our study, but we did not detect pathogens in ticks collected from either species. However, our sample size for bears was quite small (N = 2). Many of these host species, like the coyote and mule deer, maintain large home ranges, seasonal movements, and long-distance dispersal. These behaviors can increase the threat of spillover, emergence, and transmission of tick-borne disease across ecosystems [55]. Our findings are especially relevant for individuals who work outdoors or partake in recreational activities in southwest regions heavily populated by wildlife, such as border patrol agents, hikers, foresters, and hunters [58,59]. *R. parkeri* was identified in Arizona via the first human case in 2014, by 2017 seven additional cases had been reported [59].

In this study, we detected two *R. sanguineus* females with morphological abnormalities. Abnormalities in ticks can arise at any developmental stage and are thought to result from environmental stressors, such as high temperatures, use of pesticides, and pollutants on the landscape [60, 61]. Understanding tick abnormal morphologies might help to better understand a changing climate or indicate whether anomalies increase efficacy of ticks as vectors [62].

## Conclusions

Our study provides updated information on the ticks and tick-borne pathogens in southern New Mexico near the U.S.-Mexico border. Our study also helps to define the ranges of several tick species, thereby laying the foundation to detect tick range expansions as well as novel and emerging tick-borne threats. The detection of *O. megnini* and *D. albipictus* parasitizing cats was unexpected and raises concern about cats as a vehicle for moving these ticks into close contact with humans. In light of our results, sustained and expanded tick surveillance along the U.S. Mexico border is warranted.

## Acknowledgements

We thank veterinary clinics across New Mexico for supporting our research by donating ticks. Thanks to Dr. Kerry Mower for allowing us to inspect hunter harvested animals for ticks. Thanks to members of the Orr lab, Lauren MacDonald, Caitlin Curtis, Janetta Kelly and John Waller, for providing ticks from bats. Thank you to Matthew Keeling for providing ticks from a bear. We also thank Chad Kerschen, Jacob Bartley and Alana Pedersen Kamaka for their help in conducting this research.

## Supporting Information

**S1 Table. Passive Sampling**. Host species and results from passive tick sampling in New Mexico

**S2 Table. Active Sampling**. Locations and results of active tick sampling in New Mexico

